# Boosting sleep slow waves suppresses dreaming

**DOI:** 10.1101/2023.03.10.532054

**Authors:** Elsa Juan, Ceren Arslan, Franziska Regnath, Lucia M. Talamini

**Author notes:** **Corresponding author:** Lucia M. Talamini, Nieuwe Achtergracht 129 B, 1018 WS Amsterdam, The Netherlands. **Declaration of interest:** International patent application WO2018156021 of University of Amsterdam and Okazolab LtD; inventors: Talamini, Lucia Maddalena and Korjoukov, Ilia.

## Abstract

Previous findings suggest a negative correlation between slow oscillations (SO) in posterior brain regions and dreaming. Here we use a precise closed-loop auditory stimulation (CLAS) procedure to causally test whether slow oscillations suppress dreaming. Our results show that boosting posterior SO during NREM sleep decreases the likelihood of dreaming as compared to no SO boosting. This study provides the first causal evidence for the neural correlates of dreaming.

---

Sleep is characterized by alternating periods of unconsciousness and consciousness, the latter in the form of dreams. Early on, this was attributed to the alternance of NREM and REM sleep, thought to be associated, respectively, with oblivion and dreams. However, this view was revised upon consistent reporting of unconscious experiences during REM sleep (33% of REM awakenings) and dreams during NREM sleep (34% of NREM awakenings)^1,2^. Thus, considering that dreaming and unconsciousness co-exist in both REM and NREM sleep, it makes either of these sleep stages the ideal within-state model to study the neural mechanisms responsible for the modulation of consciousness during sleep.

Although REM and NREM sleep are physiologically very different, the search for the neural correlates of dreaming suggests a common mechanism across sleep stages. The most informative study, thus far, shows that in both REM and NREM sleep, dreams (in comparison to no dreams) are accompanied by a decrease in low-frequency activity (0.5-4.5 Hz) and an increase in high-frequency activity (18-25 Hz), specifically in a parieto-occipital region^3^. Other studies partially support these findings, albeit indicating that the spectral changes might be global^4^, or might include other locations, such as frontal regions^5,6^. In a study on NREM sleep, dreaming (versus not dreaming) was characterized by fewer and smaller amplitude slow waves (0.5-1 Hz) in posterior and central regions^7^. In line with these results, a TMS-EEG study showed that during NREM sleep, single-pulse TMS on the parietal cortex evoked larger negative deflection when participants were not dreaming as compared to dreaming^8^. Altogether, these correlational studies suggest that the presence of large amplitude slow oscillations – especially during NREM sleep and particularly in parietal and occipital regions – may be associated with the absence of dreams, or at least a lower level of experience^9^.

While these observations thus far are correlational, recent developments in neurostimulation methods open avenues for investigating the neural underpinnings of dreaming from a causal perspective. In particular, the presentation of short auditory stimuli at specific phases of ongoing slow oscillations, has been shown to reliably boost NREM slow wave activity, with a relatively localized effect^10–12^. So far, closed-loop acoustic stimulation (CLAS) studies have targeted frontal slow oscillations. However, using CLAS to boost slow oscillations (SO) specifically over posterior and occipital regions would constitute a unique opportunity to assess their causal role in dreaming.

In this study, we adapted a state-of-the-art CLAS procedure, based on real-time EEG modeling^13^, to target posterior slow oscillations in deep NREM sleep (sleep stage N3^14^). We recorded 26 participants (20 females; 20.2 ± 1.5 years old [mean ± SD]) randomly assigned to one of two stimulation conditions: acoustic stimulation (“STIM”; *n*=13) or sham stimulation (“SHAM”; *n*=13), during N3 sleep. In the STIM condition, our CLAS procedure presented tones precisely half-way the SO up-going slope, as targeting this phase has been shown to maximally boost slow-wave activity^11^. Targeting was directed at a posterior electrode (Pz) to specifically boost SO in the area previously associated with dreaming. The SHAM condition was identical to the STIM condition, except that no sound was played – thus no SO boosting was performed. The stimulation was continued for 10 minutes, after which participants were awoken and their dream experiences collected. Participants were then allowed to go back to sleep, and the procedure was repeated at the next occurrence of N3 until no further N3 could be detected. Participants knew they would be awoken to report their dreams, but they were blind to the stimulation conditions, and none reported hearing tones during sleep.

Across participants, a total of 3315 auditory stimuli were delivered in the STIM condition (on average 92 auditory stimuli per 10 min period) and 3033 sham stimuli in the SHAM condition (on average 91 sham stimuli per 10 min period; *t*_(24)_=0.717, *P*=0.480, *d*=0.28) (Fig 1). Overall, the targeting accuracy of our CLAS procedure was good, with only small average deviations from the target phase (0°) in both the STIM (7.68° ± 55.21°) and SHAM condition (344.81° ± 59.75°; *t*_(24)_=-0.107, *P* =0.916, *d*=-0.04).

**Fig 1.**
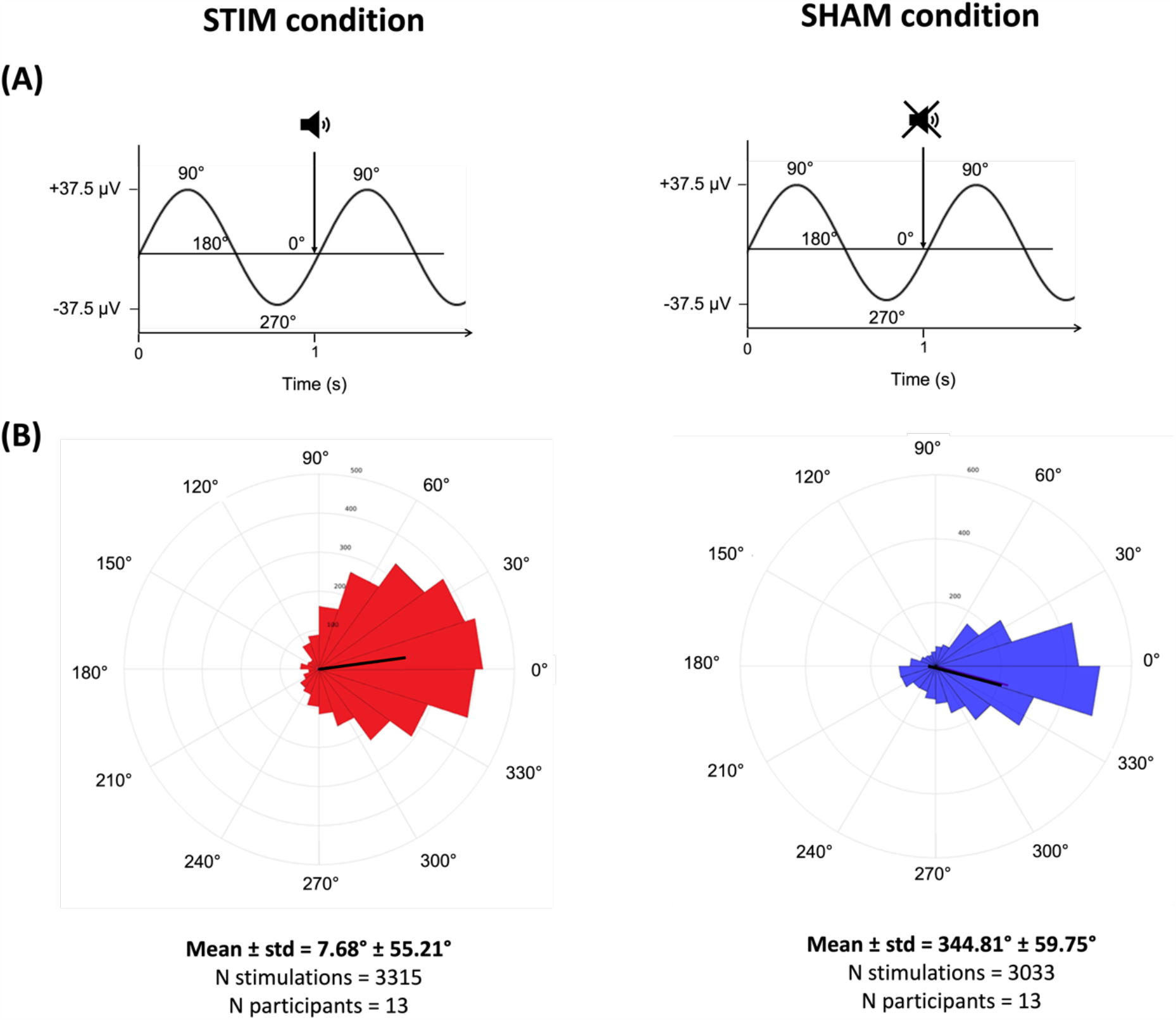
Phase prediction procedure and resulting accuracy. **(A)** The incoming signal from Pz was used to predict the phase of upcoming SOs. In both stimulation conditions, our CLAS procedure was set to target the midpoint of the SO up-going slope, defined as 0°. At this precise moment, either a 100 ms tone was released (STIM condition; left panel), or no sound was presented, but the time was marked (SHAM condition; right panel). (**B)** Across all participants, a total of 3315 auditory stimuli were released in the STIM condition and 3033 sham stimuli in the SHAM condition. The targeting accuracy was good in both conditions (respectively +8° and -15°).

In comparison to sham stimuli, auditory stimuli clearly influenced SO at Pz. Presenting a tone at 0° phase of SO (STIM condition), evoked, on average, an enhanced stimulus-locked SO-like dynamic in the seconds after stimulation. The evoked response differed significantly from SHAM between 310 to 847 ms and between 1300 to 2000 ms after stimulus onset (*P*<0.05) (Fig 2). This suggests induction and/or amplification of SO, indicating that our SO- CLAS targeting procedure successfully boosted posterior slow oscillations.

**Fig 2.**
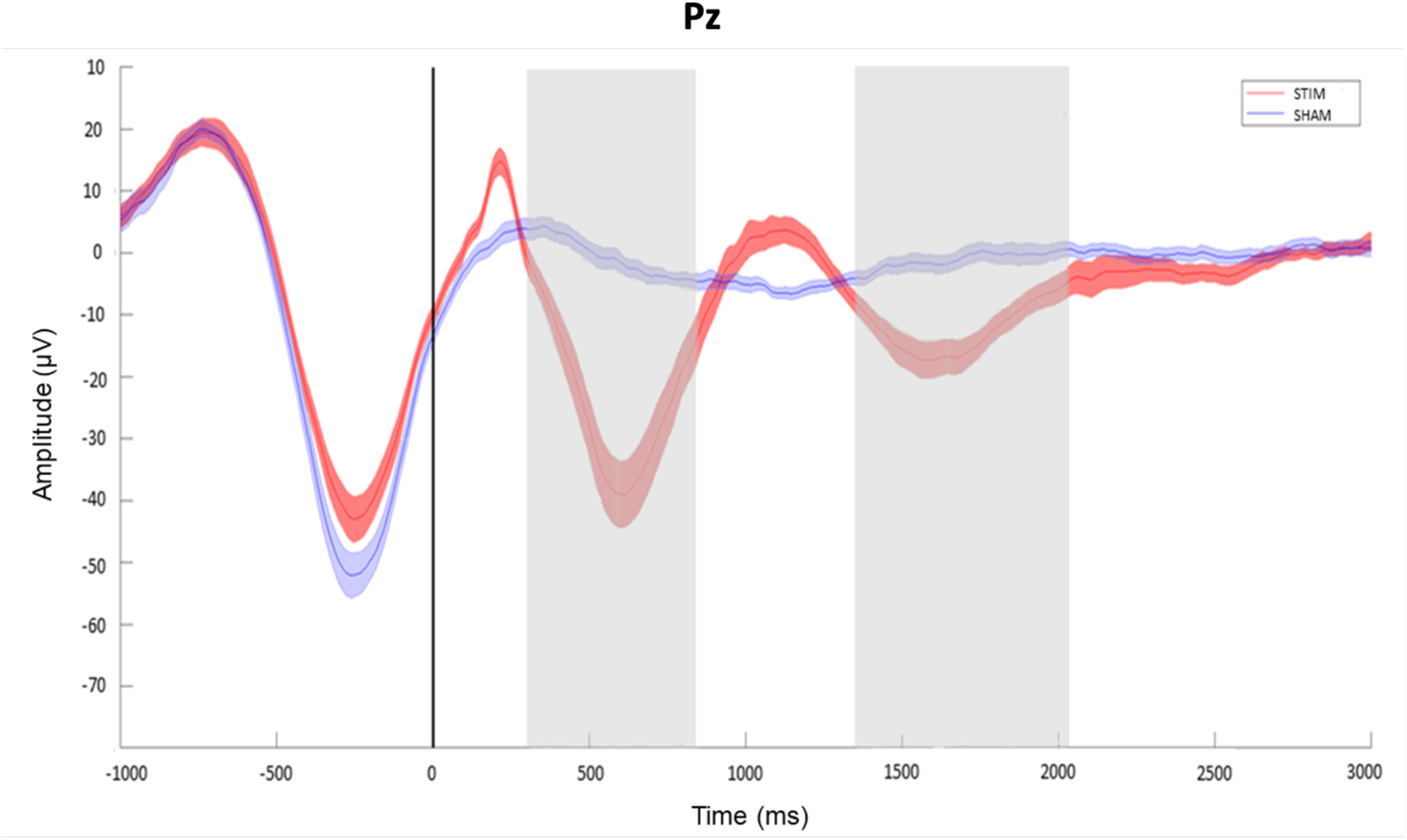
Evoked slow-wave rhythm on Pz. As compared to the SHAM condition (blue line), evoked responses were significantly boosted in the STIM condition (red line). Grey blocks indicate significant time periods; *P*<0.05). Time 0 indicates the onset of the auditory stimulus (STIM condition) or the time stamp of the sham stimulus (SHAM condition), both targeted to the 0° phase of SO on channel Pz. Red and blue shading indicate standard errors of the mean.

Crucially, our SO manipulation effectively decreased the number of dreams. Reports collected in the STIM and SHAM conditions were divided into three categories: “dreaming experiences” (DE), corresponding to any subjective experience, such as elaborate dreams or simple thoughts, “dreaming experiences without recall of content” (DEWR), when participants reported dreaming but were unable to recall the content, and “no experience” (NE), when participants reported having no conscious experience. On a total of 36 reports collected in the STIM condition, 10 (31%) were DE, 15 (46%) were DEWR and 11 (23%) were NE. In comparison, the 33 reports collected in the SHAM condition included 20 DE (59%), 9 DEWR (29%) and 4 NE (12%) (Fig 3). This shows a significant reduction of DE in the STIM condition as compared to the SHAM condition (by about 50%), along with an increase of DEWR and NE (*X*^*2*^(2,*N*=69)=7.985, *P*=0.019; Cramer’s V=0.24). Considering that both DEWR and NE indicate a degradation in consciousness and that some participants had difficulties distinguishing between these two options, we also analysed DEWR and NE together against DE. This showed an even stronger difference between conditions (*X*^*2*^(1,*N*=69)=7.55, *P*=0.006; Cramer’s V=0.33). These results indicate that boosting posterior SO caused a notable reduction of dreaming experiences and an increase of reports with partial or no conscious experience.

**Fig 3.**
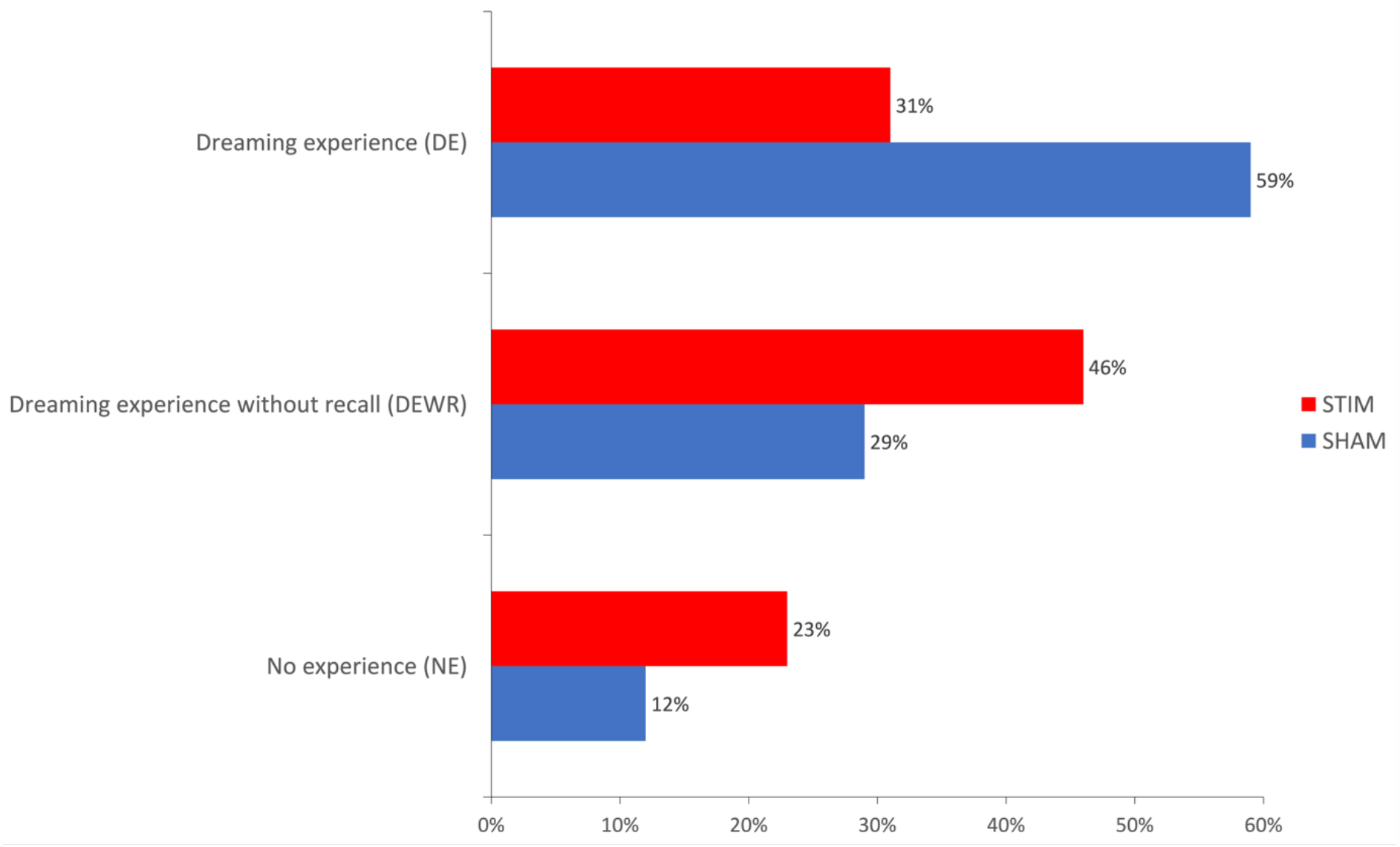
Classification of reports across stimulation conditions. As compared to spontaneous dream experiences (SHAM condition; blue bars), enhancing parietal SO (STIM condition; red bars) resulted in a decrease in dream experiences (DE) and an increase of low consciousness reports (DEWR and NE). The number of awakenings per participant was similar in both conditions (STIM: 2.8 ± 1.3 vs. SHAM: 2.5 ± 0.9; *t*_(21)_=0.53, *P*=0.601, *d*=0.21).

This study used a state-of-the-art neurostimulation technique to investigate the causal relation between slow wave activity and consciousness. We found that externally boosting slow oscillations in posterior brain regions significantly decreased the occurrence of dream experiences. Therefore, our results support the hypothesis that the presence of posterior slow oscillations is causally and negatively linked to conscious experiences during sleep.

Consistently with previous studies, our results reveal the presence of dreams during deep N3 sleep^2^, confirming that consciousness can occur during brain states dominated by low frequency activity^9^. More importantly, our study demonstrates that manipulating brain activity to increase SO dynamics in a brain area hypothesized to be associated with consciousness, leads to a decrease of dreaming experiences. Thus, it seems that while consciousness can exist against a background of EEG slow wave activity, it diminishes if SO are increased. Therefore, this study demonstrates in a clear and direct way the causal relationship between slow wave activity and consciousness. This extends previous correlational studies^3^ and brings the first, much needed, causal evidence regarding the neural underpinnings of dreaming.

Technically, we showed that posterior slow waves – which are much shallower compared to frontal ones – can nonetheless be adequately targeted using a modelling-based procedure for oscillatory phase prediction. This method is intrinsically adaptive to temporal as well as interindividual variations in the EEG signal and is thus highly suited to target a large variety of oscillations. For the purpose of this study, this state-of-the-art CLAS procedure was optimized to accurately target and effectively boost posterior SO during N3 sleep, much like reported previously for frontal SO^11,15,16^.

While we clearly established the effect of our manipulation on ongoing neural activity, our experimental set up – which capitalized on SO boosting and dream collection – was not well suited for assessing possible global sleep EEG changes accompanying the diminution of dream experience. Indeed, we adopted an established serial-awakening paradigm^2^ to collect dreams immediately after awakening, thus obtaining reliable reports and minimizing memory loss otherwise typical of dream reporting. However, the serial awakening paradigm necessarily interrupts sleep and is bound to affect overall sleep architecture variably across participants. In addition, the number of SO boosting stimuli also varied among participants, which again could affect sleep architecture. In view of these circumstances, no assessment of global sleep parameters was performed. As a further point of notice, our study is not informative regarding the specificity of posterior brain regions’ involvement in the neural correlates of dreaming. To assess this, it would be necessary to manipulate SO in other areas, such as frontal brain regions. However, given the substantial differences in EEG slow oscillation dynamics across the scalp, with much more pronounced SOs in frontal compared to more posterior areas, balancing such manipulations would not be straightforward.

This study represents the first attempt to manipulate the occurrence of dreaming experiences in real-time. This could have clinical relevance in the treatment of disorders featuring dream pathologies, like post-traumatic stress disorder (PTSD) and nightmare disorder. Furthermore, our results confirm previous findings from correlational studies, showing that SO – and particularly posterior SO – are critical for consciousness during sleep. These results provide an important step towards understanding the neural basis of dreaming.

## Online Methods

### Participants

Twenty-seven students from the University of Amsterdam, The Netherlands, participated in the study. One participant was excluded due to poor EEG signal quality. Thus, the final analyses included 26 participants [6 males, age 20.2 ± 1.5 years, 18-24 (mean ± SD, range)]. Participants were free of sleep problems, psychiatric and neurological disorders, surgical implants, cardiac problems, airway infections and epilepsy. They were asked to refrain from alcohol, cannabis, or any other drugs from 24 hours preceding the experimental night and to avoid caffeinated drinks as of 14:00. Additionally, participants were instructed to wake up no later than 08:00 and to go to bed at 23:00 for the preceding three days in order to adapt to the experimental night schedule. The study was approved by the Ethics Committee of the Department of Psychology, University of Amsterdam. Participants provided written informed consent and were compensated with course credits.

### Procedure

On the three mornings preceding their visit, participants were required to fill in the Dream Report Questionnaire (see dedicated section) to practice reporting their dreams upon awakening. On the experimental night, participants arrived at the sleep lab at 19:00. After receiving instructions, completing the informed consent form, and answering several questionnaires (not discussed here), participants were wired to a high-density EEG system. During the evening, participants performed a memory task, which is out of scope for this report and will not be discussed here. Lights were turned off at 23:00 and participants were instructed to sleep without trying to anticipate awakenings. Participants’ polysomnography signals were monitored in real time from an adjacent room. As soon as participants reached sleep stage N3, the following procedure was applied: after waiting for at least 2 minutes of consolidated N3 sleep, the experimenter turned on the CLAS procedure (see CLAS section). Half of the participants were randomly assigned to the STIM condition, in which the CLAS procedure triggered the presentation of short auditory stimuli; the other half was assigned to the SHAM condition, where no auditory stimuli were presented, but sham stimulus onset markers were recorded. After 10 minutes of continuous STIM or SHAM stimulation, participants were awoken with a long (2.5 sec) and loud (65dB) tone and prompted to answer the Dream Report Questionnaire. Questions were posed and answered verbally through an intercom and recorded to be later transcribed *verbatim*. After answering all the questions, participants were instructed to go back to sleep. This procedure was repeated for each consolidated N3 sleep segment, until around 03:00. Participants could then sleep undisturbed until 08:00. In the morning, participants received debriefing from the experimenter.

### EEG data acquisition

EEG was recorded using a 62 electrodes cap placed according to the international 10-10 system (64-channels WaveGuard original cap, ANT, Enschede, The Netherlands) with two additional electrodes placed on the earlobes as reference. In additional, bipolar electrodes were used to record horizontal eye movements (HEOG), vertical eye movements (VEOG), muscle tone (EMG) and electro-cardiography (ECG). Signals were sampled at 512 Hz using 72-channel Refa DC amplifiers and a customized version of Polybench software (TMSi, Oldenzaal, The Netherlands).

### Closed-loop acoustic stimulation (CLAS)

We applied a state-of-the-art, closed-loop acoustic stimulation (CLAS) procedure^13^ running on EventIDE software (Okazolab Ltd, London, United Kingdom) to boost posterior SO (see ^16^ for detailed description of the procedure). Specifically, oscillatory phase targeting is achieved through near real-time modelling of raw, ongoing EEG from a selected channel, through a sine fitting procedure. Model extrapolation allows prediction of the phase of incoming slow oscillations and sending auditory stimuli to coincide with the desired phase. The entire procedure is embedded in a minimal-lag, automatized, hardware-software loop. Here we selected the Pz derivation (Pz referenced to the average of both earlobe electrodes) in order to target posterior SO. In order to maximally boost SO, we set the phase parameter to present auditory stimuli to the 0° phase, corresponding to half-way the positive-going SO slope (see Fig.1). In the STIM condition, this triggered the presentation of a short (100 ms) click presented at a non-arousing volume (42dB). There was a minimum interstimulus interval of 3500 ms between clicks to ensure adequate ERP analysis. The same procedure was followed in the SHAM condition, but no clicks were presented. In case there was any sign of arousal during the stimulation period, the procedure was immediately interrupted; it was resumed upon detection of a new consolidated N3 period, with the volume reduced to 40 dB.

### Dream report questionnaire

On average, the first awakening was done at 00:20 and the last awakening at 02:30. The dream report questionnaire (adapted from^2^, available upon reasonable request) was administered at each awakening to collect participants’ subjective experiences. The duration of the interview lasted between two and nine minutes (mean±SD: 5±2 min), depending on how many questions the participant had to answer. The first question was always an open question aimed at collecting a free report of participant’s experience: “*What was the last thing going through your mind prior to the alarm sound*?”. As in Siclari et al. (2013)^2^, participants were instructed to report only their most recent subjective experience and not the full experience they might have had. Based on this question, participant’s response was classified in three categories (1) “Dreaming experience” (DE), meaning that the participant reported some type of mental activity, including dreams but also simple thoughts or sensations; (2) **“**Dreaming experience without recall” (DEWR) when the participant felt like they had a dream but couldn’t remember the content; (3) “No experience” (NE) if the participant was certain there was nothing going on in their mind right before the alarm. Following DE reports, twenty-one questions regarding the emotional valence, arousal level and duration of the subjective experience were administered; for DEWR and NE, a subset of six questions was collected. At the end of the questionnaire, participants were told to go back to sleep.

### Behavioral analyses

A chi-square test of independence was run to test if the distribution of answers across categories (DE, DEWR and NE) differed by condition. Statistical analyses were run using JASP^17^.

### Oscillatory phase targeting performance

To check the accuracy of the CLAS procedure, we calculated the phase targeting accuracy of all delivered auditory and sham stimuli. Signal from channel Pz was filtered between 0.5 – 1.5 Hz using a zero-phase shift Butterworth filter. The phase corresponding to the onset of each stimulus was computed using Hilbert transform. The phase distribution for each condition was then examined using the CircStat toolbox^18^.

### Pre-processing

EEG data was pre-processed using the EEGLAB v2020 toolbox^19^ for MATLAB 2019b (Mathworks, Natrick, MA). Data was filtered between 0.1 – 45 Hz and re-referenced to the average of the signals of the two earlobe electrodes. The continuous data from the last 10 min before each awakening for each condition was concatenated and considered for analyses. We visually inspected the EEG data to reject epochs with arousals and muscle artifacts. EEG channels with poor signal were removed and interpolated from neighbouring channels using spherical splines (NetStation, Electrical Geodesic Inc.).

### Event-related potentials (ERPs)

The signal from channel Pz (referenced to the average of the two earlobe electrodes) was epoched between -1000 and 3000 ms. The Monte Carlo sampling method with cluster correction^20^ was used to evaluate possible ERP differences between conditions.

## Acknowledgements

We would like to thank Ilia Korjoukov for his assistance with EventIDE software (OkazoLab, 2019).

## Data availability

The data is available upon reasonable request.

## References

1. Nielsen, T. A. A review of mentation in REM and NREM sleep: ‘Covert’ REM sleep as a possible reconciliation of two opposing models. Behav. Brain Sci. 23, 851–866 (2000).

2. Siclari, F., Larocque, J. J., Postle, B. R., Tononi, G. & Windt, J. M. Assessing sleep consciousness within subjects using a serial awakening paradigm. 4, 1–9 (2013).

3. Siclari, F. et al. The neural correlates of dreaming. Nat. Neurosci. 20, 872–878 (2017).

4. Zhang, J. & Wamsley, E. J. EEG predictors of dreaming outside of REM sleep. Psychophysiology 56, 1–14 (2019).

5. Chellappa, S. L., Frey, S., Knoblauch, V. & Cajochen, C. Cortical activation patterns herald successful dream recall after NREM and REM sleep. Biol. Psychol. 87, 251–256 (2011).

6. Scarpelli, S. et al. Predicting Dream Recall: EEG Activation During NREM Sleep or Shared Mechanisms with Wakefulness? Brain Topogr. 30, 629–638 (2017).

7. Siclari, F., Bernardi, G., Cataldi, J. & Tononi, G. Dreaming in NREM sleep: A high-density EEG study of slow waves and spindles. J. Neurosci. 38, 9175–9185 (2018).

8. Nieminen, J. O. et al. Consciousness and cortical responsiveness: A within-state study during non-rapid eye movement sleep. Sci. Rep. 6, 1–10 (2016).

9. Frohlich, J., Toker, D. & Monti, M. M. Consciousness among delta waves: a paradox? Brain (2021) doi:10.1093/brain/awab095.

10. Talamini, L. M. Memory Manipulation During Sleep: Fundamental Advances and Possibilities for Application. 313–334 (2017) doi:10.1007/978-3-319-45066-7_19.

11. Cox, R., Korjoukov, I., De Boer, M. & Talamini, L. M. Sound asleep: Processing and retention of slow oscillation phase-targeted stimuli. PLoS One 9, (2014).

12. Ngo, H. V. V., Claussen, J. C., Born, J. & Mölle, M. Induction of slow oscillations by rhythmic acoustic stimulation. J. Sleep Res. 22, 22–31 (2013).

13. Talamini, L. M. & Korjoukov, I. Assembly, method and computer program product for influencing a biological process. (2018).

14. Iber, C., Ancoli-Israel, S., Chesson, A. L. & Quan, S. F. The new sleep scoring manual - The evidence behind the rules. J. Clin. Sleep Med. 3, 107 (2007).

15. Ngo, H. V. V., Martinetz, T., Born, J. & Mölle, M. Auditory closed-loop stimulation of the sleep slow oscillation enhances memory. Neuron 78, 545–553 (2013).

16. Pathak, V., Juan, E., van der Goot, R. & Talamini, L. M. The looping lullaby: closed-loop neurostimulation decreases sleepers’ sensitivity to environmental noise. IEEE Control Syst. Mag. 2009–2011 (2021).

17. Love, J. et al. JASP: Graphical statistical software for common statistical designs. J. Stat. Softw. 88, (2019).

18. Berens, P. CircStat: A MATLAB Toolbox for Circular Statistics. J. Stat. Softw. 31, 1–21 (2009).

19. Delorme, A. & Makeig, S. EEGLAB: an open source toolbox for analysis of single-trial EEG dynamics including independent component analysis. J. Neurosci. Methods 134, 9–21 (2004).

20. Maris, E. & Oostenveld, R. Nonparametric statistical testing of EEG- and MEG-data. J. Neurosci. Methods 164, 177–190 (2007).

